# HIV-1 CD4-binding site germline antibody–Env structures inform vaccine design

**DOI:** 10.1101/2022.03.25.485873

**Authors:** Kim-Marie A. Dam, Christopher O. Barnes, Harry B. Gristick, Till Schoofs, Michel C. Nussenzweig, Pamela J. Bjorkman

## Abstract

BG24, a VRC01-class broadly neutralizing antibody (bNAb) against HIV-1 Env with relatively few somatic hypermutations (SHMs), represents a promising target for vaccine strategies to elicit CD4-binding site (CD4bs) bNAbs. To understand how SHMs correlate with BG24 neutralization of HIV-1, we solved 4.1 Å and 3.4 Å single-particle cryo-EM structures of two inferred germline (iGL) BG24 precursors complexed with engineered Env-based immunogens lacking CD4bs N-glycans. Structures revealed critical Env contacts by BG24_iGL_ and identified antibody light chain structural features that impede Env recognition. In addition, biochemical data and cryo-EM structures of BG24_iGL_ variants bound to Envs with CD4bs glycans present provided insights into N-glycan accommodation, including structural modes of light chain adaptations in the presence of the N276_gp120_ glycan. Together, these findings revealed Env regions critical for germline antibody recognition and potential sites to alter in immunogen design.

---

Current strategies to engineer a vaccine towards preventing HIV-1 infection involve designing Env-mimetic immunogens that can elicit broadly neutralizing antibodies (bNAbs)^1–4^. The CD4-binding site (CD4bs) epitope is a target of immunogen design as bNAbs in this class have been shown to be among the most potent and broad^5–9^. Several studies have shown passive immunization using CD4bs bNAbs can confer protection from HIV-1 infection in animal models and human clinical trials, suggesting that immunization strategies to elicit these antibodies at effective concentrations would also be protective^6, 10–17^. This includes the VRC01-class of bNAbs that are derived from the VH1-2*02 variable heavy chain gene segment and are characterized by a short 5 amino acid complementary determining region 3 (CDRL3) in the antibody (Ab) light chain and a shortened or flexible CDRL1^5, 18^. These characteristics are necessary for VRC01-class bNAbs to accommodate the heavily N-glycosylated landscape of the CD4bs of HIV-1 Envs. Thus, VRC01-class bNAbs generally require high levels of somatic hypermutation (SHM), which is challenging to elicit through vaccination.

Germline precursors of bNAbs do not generally show detectable binding to HIV-1 Envs^19, 20^, therefore, the germline targeting approach to HIV-1 vaccine design involves efforts to engineer immunogens that can engage germline B cell receptors (BCRs) and initiate bNAb development^21^. Inferred germline (iGL) versions of mature bNAbs derived from predicted germline gene segment sequences represented in the human B cell repertoire^22, 23^ are used for the germline targeting approach. Analysis of VRC01-class iGLs has shown that the human VH1-2*02 heavy chain gene segment encodes signature residues that are required for breadth and potency^18^. Furthermore, germline VRC01-class precursors have been isolated from naïve individuals and mature bNAbs have been identified from multiple HIV-1-infected human donors, suggesting that raising this class of bNAbs is not uncommon in natural infection^24, 25^. Taken together, VRC01-class bNAbs are attractive targets for immunogen design.

The VRC01-class of bNAbs target the CD4bs, a particularly challenging epitope to elicit bNAbs against due to the presence of the CD4bs N-glycans that sterically obstruct interactions between Env and Ab CDRs^26^. The glycan at position N276_gp120_ is highly conserved and poses the greatest steric barrier to binding VRC01-class bNAb iGLs, as Ab residues in the iGL CDRL1 that interact with this region are typically 11-12 residues and cannot accommodate the N276_gp120_ glycan. Mature CD4bs Abs develop shortened or flexible CDRL1s to accommodate this glycan^24, 27, 28^. Thus, understanding the structural basis for how CD4bs iGL Abs mature to effectively accommodate the N276_gp120_ glycan is essential in efforts to develop effective immunogens to prime VRC01-class iGL precursors and shepherd antibody responses towards bNAb development. Furthermore, an overall structural understanding of VRC01-class iGL recognition of HIV-1 Envs and immunogens is limited as the only existing Fab-Env structures involving germline CD4bs Abs are complexed with gp120 or Env trimer immunogens lacking the N276_gp120_ glycan^3, 23, 29^. In addition, in the case of an iGL Fab complexed with an Env trimer, obtaining a structure required chemical cross-linking between the Env and Ab to form a stable complex^22^.

A new VRC01-class bNAb isolated from an elite neutralizer, BG24^30^, is an attractive target for germline-targeting immunogen design. BG24 shows similar neutralization and breadth to other CD4bs bNAbs, but includes only 22.6% and 19.5% amino acid substitution by SHM in variable heavy and light chain genes, respectively^30^, as compared with higher levels of amino acid substitution in VRC01-class bNAbs^7, 9, 28, 31^, with the exception of the PCIN63 lineage that has similar levels of SHM to BG24^32^. Structural characterization of BG24 bound to the clade A BG505 Env revealed a similar binding orientation to more mutated VRC01-class bNAbs, and signature contacts common to VRC01-class bNAbs ^30^. Furthermore, neutralization studies using variants of BG24 that reverted variable heavy (V_H_) and variable light (V_L_) domain residues to germline counterparts showed that even fewer SHMs were necessary to maintain neutralization breadth^30^. Collectively, this suggests broad and potent neutralization targeting the CD4bs could be achieved through immunization without stimulating high levels of SHM.

To better understand how the BG24 bNAb was elicited and inform VRC01-class immunogen design, we structurally characterized the binding of two versions of the BG24 iGL to the CD4bs germline-targeting immunogen BG505-SOSIPv4.1-GT1^3^ (hereafter referred to as GT1). We solved two single-particle cryo-electron microscopy (cryo-EM) structures of GT1 in complex with BG24_iGL_s containing either mature or iGL CDR3s at 4.1 Å and 3.4 Å resolution, respectively, in both cases in the absence of chemical cross-linking. Furthermore, to understand how N-glycans impact germline Ab recognition of Env, we conducted biochemical assays and solved cryo-EM structures of BG24_iGL_ derivatives bound to Envs that included the N276_gp120_ glycan. The structures demonstrated that the CDRL1s of BG24_iGL_s can adopt conformations that accommodate the N276_gp120_ glycan, an important capability for a germline-targeting CD4bs immunogen. Collectively, these structures provide information regarding the physical characteristics of iGLs that recognize HIV-1 Env and provide a structural basis for the design of immunogens engineered to engage and mature germline Abs.

## Results

### Cryo-EM structures of GT1-BG24_iGL_-101074 complexes

To gain insight into how BG24 precursors interact with an HIV-1 Env-based immunogen, we created iGL versions of BG24 and used single-particle cryo-EM to structurally characterize them in complex with GT1, a CD4bs germline-targeting immunogen^3^. GT1 was modified from a soluble clade A BG505 SOSIP.664 native-like Env trimer^33^ to permit binding of VRC01-class germline precursors by including T278R_gp120_ and G271S_gp120_ substitutions and mutations to remove potential N-linked glycosylation sites (PNGSs) at positions N276_gp120_, N462_gp120_, N386_gp120_, and N197_gp120_ in the CD4bs^3^. Two iGL versions of BG24 Fab constructs were made starting with the VH1-2*02 and VL2-11*01 heavy and light chain germline gene segment sequences: one containing the CDR3s from mature BG24 (BG24_iGL-CDR3mat_) and the other containing the iGL CDR3s (BG24_iGL-CDR3iGL_) (Fig. 1a). Each BG24_iGL_ was structurally characterized in complex with GT1 and the V3 bNAb 10-1074^34^.

**Fig. 1.**
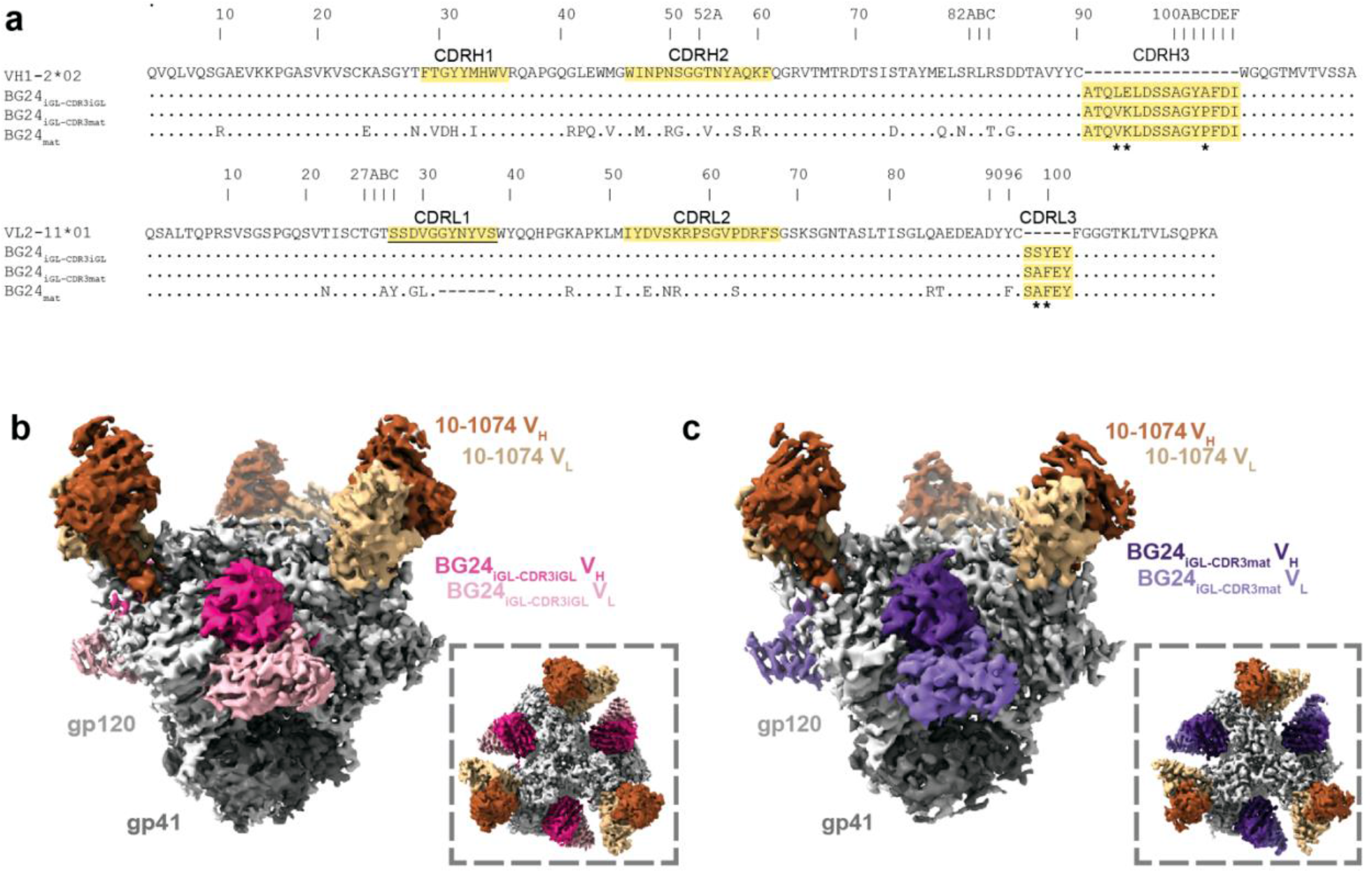
BG24_iGL_s bind the CD4bs of the GT1 immunogen. **a**, Sequence alignment of V_H_ and V_L_ iGL precursors of BG24 (VH1_2*02 and VL2_11*01), BG24_iGL-CDR3iGL_, BG24_iGL-CDR3mat_, and BG24_mat_. CDRs are highlighted in yellow. Asterisks (*) indicate residue differences between mature and iGL CDR3s. Underlined CDRL1 indicates sequence used for the CDRL1 in the BG24_CDRL1-iGL_ construct. **b**,**c,** Side and top-down (inset) views of cryo-EM density of BG24_iGL-CDR3iGL_-GT1-10-1074 (**b**) and BG24_iGL-CDR3mat_-GT1-10-1074 (**c**).

Cryo-EM structures of BG24_iGL-CDR3iGL_ and BG24_iGL-CDR3mat_ Fabs bound to GT1 were solved at 3.4 Å and 4.1 Å, respectively (Fig. 1b,c, Extended Data Fig. 1a-j, Supplementary Table 1). Both 3D cryo-EM reconstructions showed three BG24_iGL_ and three 10-1074 Fabs bound per Env trimer. However, for the BG24_iGL-CDR3iGL_-GT1-101074 complex, a distinct 3D class contained two BG24-iGL_CDR3iGL_ Fabs bound to the GT1 Env (Extended Data Fig. 1e-f, Supplementary Table 1). We also solved a 1.4 Å crystal structure of unbound BG24_iGL-CDR3mat_ Fab (Extended Data Fig. 1k, Supplementary Table 2), which exhibited six disordered residues within CDRL1, but otherwise superimposed with a 1.3 Å root mean square deviation (rmsd; calculated for 225 V_H_-V_L_ Cα atoms) with the Env-bound BG24_iGL-CDR3mat_ Fab structure, suggesting no major structural differences upon Env binding.

### BG24_iGL_ Fabs recognize the modified CD4bs in GT1 Env

The GT1 complexes with BG24_iGL_s included density for CD4bs N-glycans attached to residues N234_gp120_, N363_gp120_, and N392_gp120_ (Fig. 2a-b). These N-glycans were also observed in the crystal structure of BG505 Env complexed with a mature BG24 Fab^30^ (BG24_mat_) (PDB 7UCE), which also included densities for N-glycans at N197_gp120_, N276_gp120_, and N386_gp120_ that are not present in GT1 (Fig. 2c). Despite additional glycans in BG505 compared with GT1, the CDR loops in the GT1-bound iGL Fabs showed similar orientations and positions as in the BG505-bound BG24_mat_ Fab, except for CDRL1, which is six residues longer in BG24_iGL_ than in BG24_mat_ (Fig. 1a, 2d-f).

**Fig. 2.**
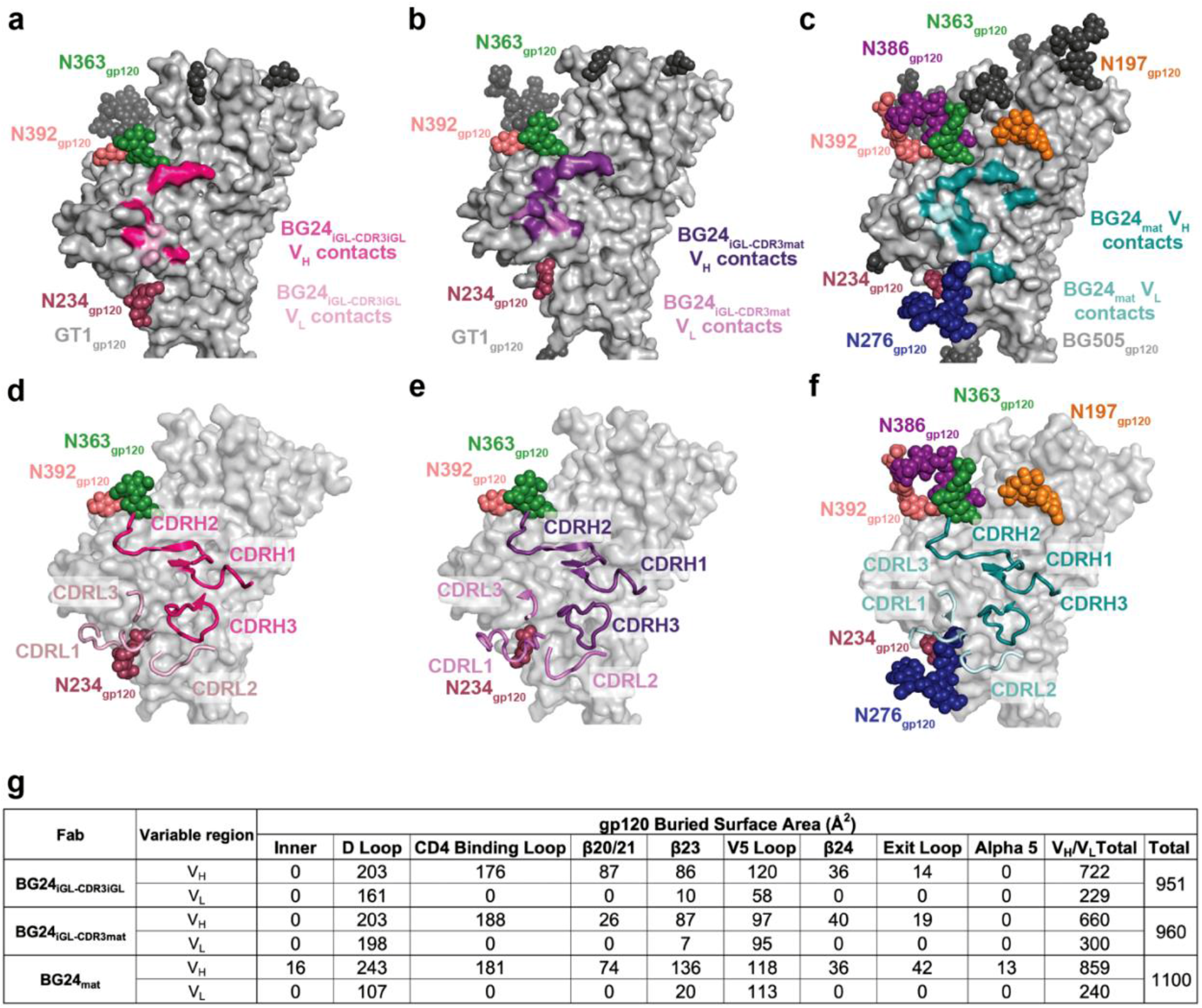
Comparison of BG24_iGL_ and BG24_mat_ CD4bs epitopes. Surface contacts made by BG24_iGL-CDR3iGL_ V_H_ (dark pink) and V_L_ (light pink) on GT1 gp120 (**a**), BG24_iGL-CDR3mat_ V_H_ (dark purple) and V_L_ (light purple) on GT1 gp120 (**b**), BG24_mat_ V_H_ (deep teal) and V_L_ (light teal) surface contacts on BG505 by BG24_mat_ (PDB 7UCF) (**c**). Surface representation of gp120 with cartoon representations of BG24_iGL-CDR3mat_ (**d**), BG24_iGL-CDR3iGL_ (**e**), and BG24_mat_ (**f**) CDR loops. **g**, Summary table of gp120 buried surface area (BSA) (Å_2_) calculations for BG24_iGL-CDR3iGL_, BG24_iGL-CDR3mat_, and BG24_mat_ at the inner domain (inner), D loop, CD4bs loop, β20/21, β23, V5 loop, β24, exit loop, and alpha 5 regions of the CD4bs. BSA calculations were conducted for gp120 peptide components and did not include glycan interactions.

BG24_iGL-CDR3mat_ and BG24_iGL-CDR3iGL_ buried comparable surface areas on GT1 gp120 (960 Å^2^ and 951 Å^2^, respectively) as compared with an only slightly larger surface area (1099 Å^2^) buried on BG505 gp120 in the BG24_mat_-BG505 structure (PDB 7UCF) (Fig. 2a-b, g). However, differences in interactions between the BG24_iGL_-GT1 and BG24_mat_-BG505 structures suggest that SHM substitutions enrich interactions in particular regions within the CD4bs. For example, in the BG24_mat_-BG505 complex, BG24_mat_ residue S99_HC_ hydrogen bonds with the gp120 inner domain residue K97_gp120_ (Fig. 2c, 2g). K97_gp120_ is ∼90% conserved among HIV-1 Envs, making this a crucial interaction of broad and potent CD4bs bNAbs^18^. Residue S99_HC_ is a germline encoded residue, however, in both BG24_iGL_-GT1 structures, it did not hydrogen bond with K97_gp120._ Compared to BG24_iGL_-GT1, BG24_mat_-BG505 also showed increased V_H_ buried surface area (BSA) in the gp120 exit loop (gp120 residues 470-474), and gained BSA in the gp120 alpha 5 region, suggesting BG24_mat_ has a larger BSA footprint than the BG24_iGL_s in these CD4bs regions (Fig. 2c, 2g). Interestingly, the overall interface BSA values for the gp120 peptide components of the BG24_iGL_-GT1 and BG24_mat_-BG505 structures were similar (960 and 951 Å^2^ vs 1099 Å^2^, respectively). Although a germline precursor antibody presumably exhibits fewer contacts to an antigen than its counterpart somatically-mutated bNAb, the modifications in GT1 (both amino acid substitutions and removals of N-glycans) allowed increased contacts between BG24_iGL_s and the GT1 gp120.

### BG24 somatic hypermutation plays a role in CD4bs recognition

We next compared how differences in BG24_iGL_ and BG24_mat_ contribute to their recognition of GT1 and BG505, respectively. BG24_iGL_ contains a germline 11-residue CDRL1 that can recognize the mostly aglycosylated CD4bs in GT1, whereas the BG24_mat_ CDRL1 is six residues shorter and includes a glycine to create a more flexible loop that can accommodate the N276_gp120_ glycan^30^. In the BG24_mat_-BG505 structure, the five-residue BG24 CDRL1 is oriented adjacent to the N276_gp120_ glycan (Fig. 3a). The CDRL1 interface with GT1 in the BG24_iGL-CDR3iGL_ and BG24_iGL-CDR3mat_ structures showed the longer CDRL1s in the germline precursor V_L_ domains in different conformations, demonstrating CDRL1 flexibility (Fig. 3b-c) consistent with cryo-EM data processing. Indeed, the local resolutions for the CDRL1 in these structures were poor and resolved only after iterative rounds of focused classification and local refinements (Extended Data Fig. 2a-b). Overlaying the BG24_iGL_ CDRL1s with the gp120 region surrounding the N276_gp120_ glycan from the BG24_mat_-BG505 structure showed steric clashes, consistent with SHM being necessary for N276_gp120_ glycan accommodation by BG24 (Fig. 3b-c).

**Fig. 3.**
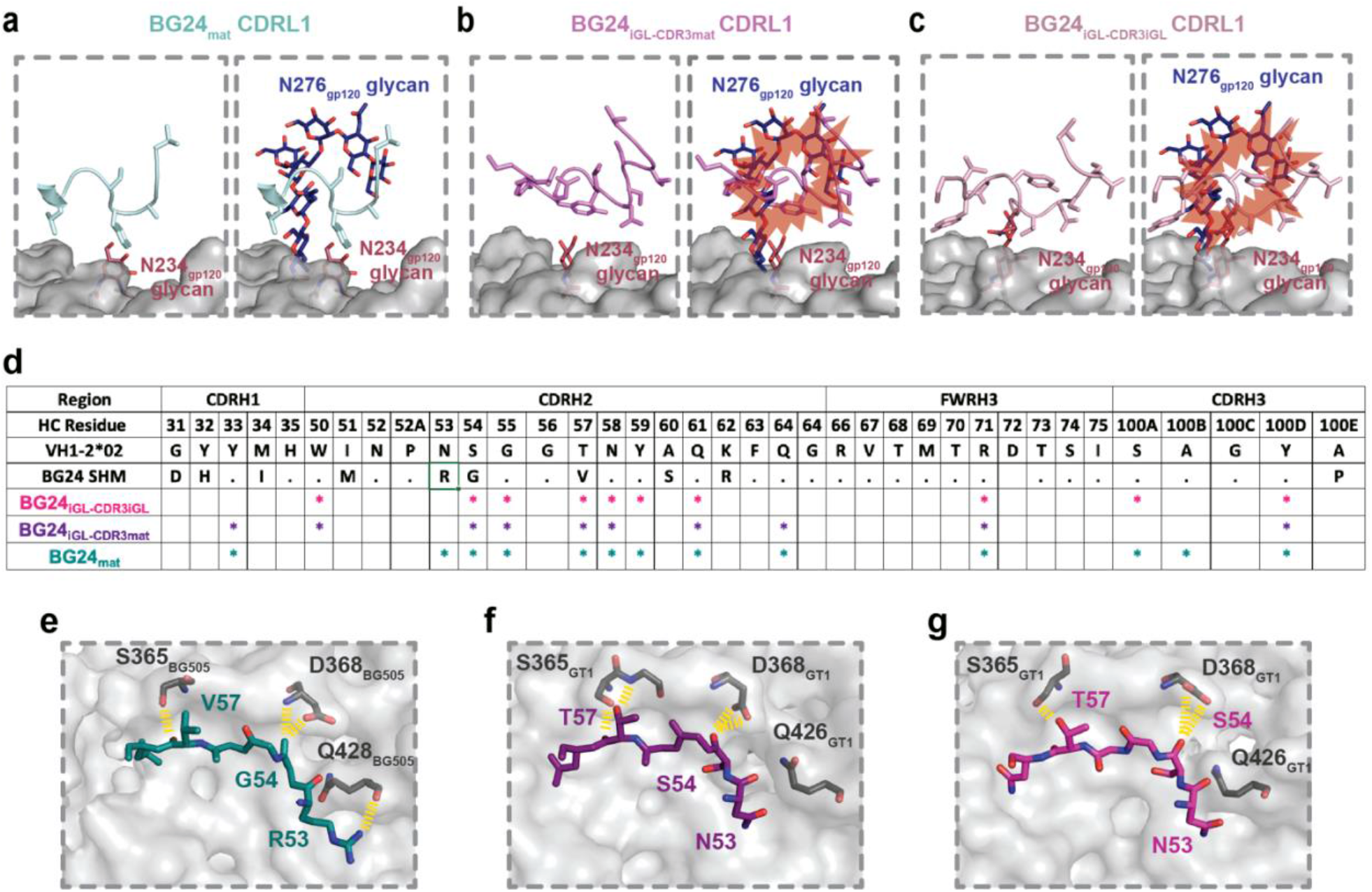
Somatic hypermutation plays a role in BG24 recognition of the CD4bs interface. gp120 surface in the vicinity of the CD4bs with cartoon representation main chain/stick side chains for the CDRL1s of **a**, BG24_mat_, **b**, BG24_iGL-CDR3mat_, **c**, BG24_iGL-CDR3iGL_ overlaid with the N276_gp120_ N-glycan from the BG24_mat_-BG505 complex (PDB 7UCF). Steric clashes are represented with red bursts. **d,** Table summarizing HC paratope residues in BG24_iGL-CDR3iGL_-GT1, BG24_iGL-CDR3mat_-GT1 and BG24_mat_-BG505 structures. The paratope was defined by Ab residues that make contacts with gp120 within 4 Å for each structure. Stick representations of the CDRH2 residues from **e**, BG24_mat_, **f**, BG24_iGL-CDR3mat_, **g**, BG24_iGL-CDR3iGL_ interacting with BG505 or GT1 gp120 residues. Yellow dashed lines indicate Ab-gp120 interactions within 4 Å.

The role of SHMs in Env recognition is summarized in Fig. 3d, where BG24_iGL-CDR3iGL_, BG24_iGL-CDR3mat_, and BG24_mat_ HC paratope interactions are mapped to individual Ab residues within 4 Å of gp120. Paratope contacts were limited to CDRs H1, H2, and H3, as well as framework region 3 in the heavy chain (FWRH3), with most contacts in CDRH2. Previous studies showed neutralization by an engineered BG24 “minimal” construct that contained germline-reverted SHMs in FWRs, CDRH1, and CDRL2, but maintained most SHMs in CDRH2, suggesting the importance of SHMs in this region^30^. The structure of BG24_mat_-BG505 showed a CDRH2 SHM (N53R_HC_) interacted with Q428_gp120_ in gp120 β20/21 (Fig. 1a, 3e). β20/21 interactions with germline-encoded N53_HC_ were absent in BG24_iGL-CDR3iGL_-GT1 and BG24_iGL-CDR3mat_-GT1 (Fig. 3f-g). This demonstrates the direct impact of SHM in creating favorable interactions with Env. Other BG24_mat_ somatically hypermutated residues in CDRH2 also interacted with the CD4bs loop (gp120 residues 356-371); e.g., residue T57V_HC_ makes a backbone interaction with S365_gp120_, and S54G_HC_ interacts with D368_gp120_, a highly conserved Env residue (Fig. 3d)^18^ For BG24_iGL_, germline-encoded residues at positions T57_HC_ and S54_HC_, maintain similar interactions with GT1 residues S365_gp120_ and D368_gp120_, respectively, suggesting that these germline encoded residues may be sufficient to engage the CD4bs loop (Fig. 3f-g).

Signature residues encoded by the VH1-2*02 germline gene in VRC01-class bNAbs interact with conserved gp120 residues and are correlated with neutralization potency^18^. These interactions have been structurally characterized in the context of VRC01-class iGLs bound to monomeric gp120s^22, 23, 29^, but there are no known structures of VRC01-class iGLs bound to a trimeric Env, except when the iGL was chemically cross-linked to Env^22^. To evaluate and verify VRC01-class VH1-2*02 germline-encoded interactions with an Env trimer, we compared these interactions in the BG24_iGL_-GT1 and BG24_mat_-BG505 structures (Extended Data Fig. 2c-f). Specifically, as previously described in structures involving gp120 monomers^22, 23, 29^, germline-encoded R71_HC_ in the BG24_iGL_-GT1 and BG24mat-BG505 structures formed a salt bridge with the conserved D368_gp120_ side chain, an Ab interaction that mimics the interaction of host receptor residue R59_CD4_ with D368_gp120_ (Extended Data Fig. 2c). In the gp120 D loop, there were interactions between the backbone and side chain of N280_gp120_ with Y100B_HC_ and germline-encoded W50_HC_ side chains (Extended Data Fig. 2d). In the V5 loop, interactions between the conserved R456_gp120_ residue and germline-encoded N58_HC_ are conserved in both structures (Extended Data Fig. 2e). In BG24_iGL CDR3mat_-GT1, atoms within these Fab residues were separated by more than 5 Å from atoms within gp120 residues; thus, this is not defined as an interaction. In the light chain, E96_LC_ interacted with the backbone of G456_gp120_ and the sidechain of N280_gp120_ (Extended Data Fig. 2f).

### GT1 CD4bs glycan modifications affect BG24 binding

To evaluate how glycan modifications in the GT1 immunogen contributed to BG24_iGL_ binding, we evaluated the binding of BG24 constructs to GT1 with Env PNGSs either restored to or removed from the CD4bs (Fig. 4a). The BG24 constructs included BG24_mat_, BG24 with germline CDRL1 (BG24_CDRL1-iGL_) (Fig. 1a), BG24_iGL-CDR3mat_, BG24_iGL-CDR3iGL_, and BG24 with an iGL light chain (BG24_LC-iGL_). PNGSs were individually restored at positions N197_gp120_, N276_gp120_, N386_gp120_, and N462_gp120_ and removed at N234_gp120_ to create five GT1 constructs with altered glycan landscapes (GT1_N197gp120_, GT1_N276gp120_, GT1_N386gp120_, GT1_N462gp120_, and GT1_del N234gp120_, respectively). BG505 and GT1 binding was evaluated by enzyme-linked immunosorbent assays (ELISAs). Restoring Env PNGSs at positions N197_gp120_, N386_gp120_, and N462_gp120_ and removing the PNGS at N234_gp120_ did not greatly affect binding of BG24 IgG constructs (Fig. 4a). BG24_iGL_ constructs did not bind detectably to GT1 with a PNGS at N276_gp120_; however, BG24_mat_ and mature BG24 constructs with iGL LC features (BG24_CDRL1-iGL_, BG24_LC-iGL_) showed comparable binding to each other on GT1_N276gp120_ (Fig 4a).” BG24_mat_ was the only Ab that showed substantial binding to BG505, which unlike the GT1 Env, included all PNGSs. We conclude that BG24 constructs with a long, germline CDRL1 can accommodate the N276_gp120_ glycan on Envs that have been engineered to have a limited glycan landscape in the CD4bs.

**Fig. 4.**
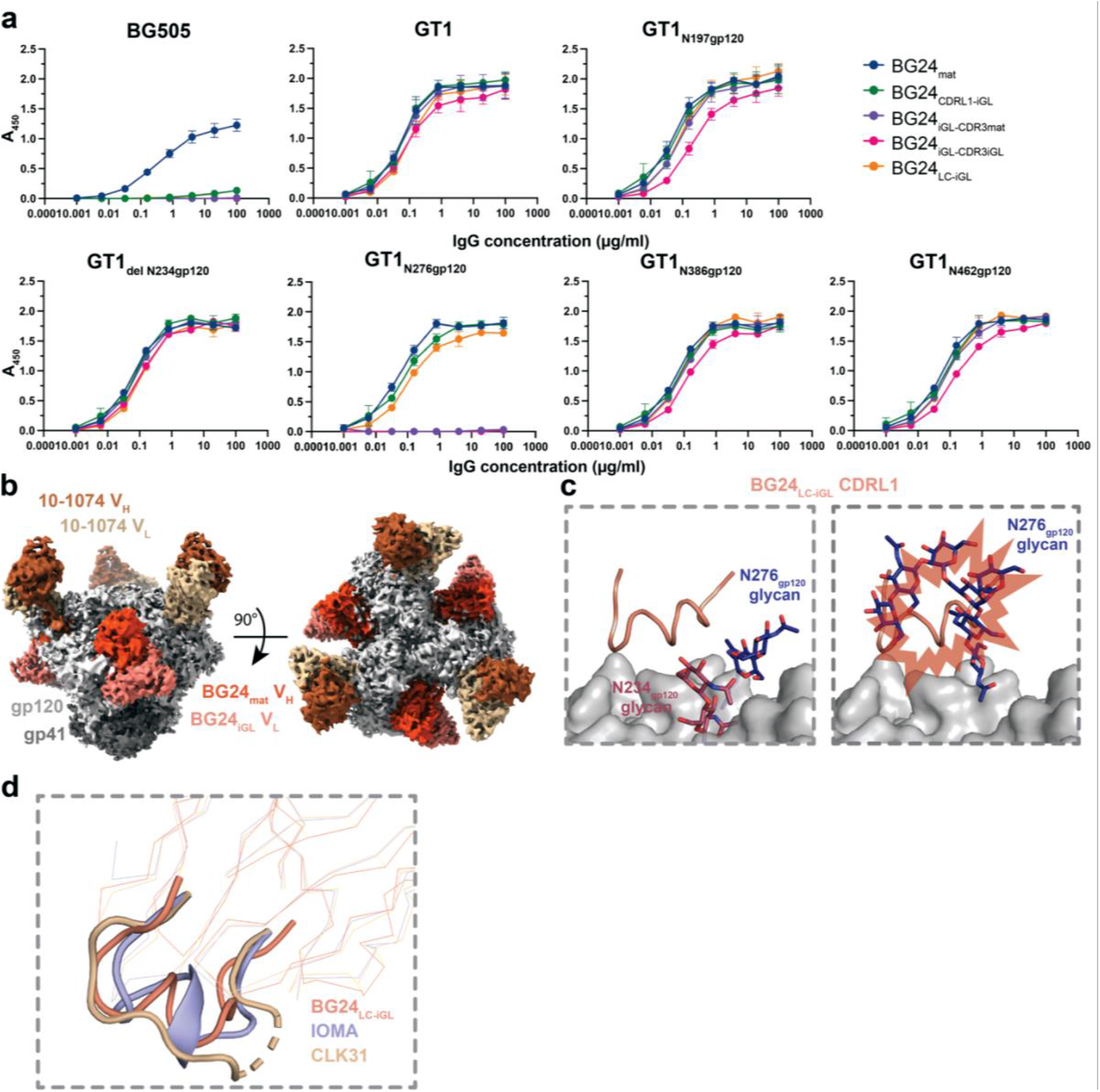
BG24_iGL_ binding is mediated by CD4bs glycans. **a**, ELISA to access binding of the indicated BG24 Abs to BG505, GT1, and GT1 SOSIP Envs with altered N-glycans in the CD4bs. Streptavidin plates were coated with randomly biotinylated SOSIPs and incubated with IgGs at increasing concentrations. Values are shown as mean of two individual replicates with associated error bars. **b**, Side and top-down views of cryo-EM density of BG24_LC-iGL_-GT1-10-1074. **c**, Cartoon representation of the CDLR1 of BG24_LC-iGL_ (left) and overlaid with the N276_gp120_ N-glycan from a BG24_mat_-BG505 (PDB 7UCF) (right). Predicted steric clashes are indicated by red bursts. **d,** Alignment of BG24_LC-iGL_ (from the BG24_LC-iGL_-GT1_N276gp120_-10-1074 structure), IOMA (PDB 5T3Z), and CLK31 (PDB 6P2P) LC with CDRL1s represented in cartoon.

To gain insight into BG24 CDRL1-iGL interactions in GT1 with an N276_gp120_ glycan, we solved a single-particle cryo-EM structure of BG24_LC-iGL_ bound to GT1 containing the restored N276_gp120_ PNGS (GT1_N276gp120_) (Fig. 4b, Extended Data Fig. 3a-h, Supplementary Table 1). We identified three unique 3D volumes containing either one, two, or three bound BG24_LC-iGL_ Fabs, with the highest resolution complex (4.2 Å) being C3 symmetric with three bound BG24_LC-iGL_ Fabs (Extended Data Fig. 3a-h, Supplementary Table 1). Electron density in the Fab CDRL1 was not optimal after cryo-EM processing; therefore, side chains were not modeled (Extended Data Fig. 3i).

The BG24_LC-iGL_-GT1_N276gp120_ complex structure showed that the Fab CDRL1 main chain residues adopted a helix-like conformation to accommodate the N276_gp120_ glycan (Fig. 4c). Available crystallographic and cryo-EM Env structures demonstrate that the N276_gp120_ glycan is conformationally heterogeneous (Extended Data Fig. 3h). Indeed, the N276_gp120_ glycans in the GT1_N276gp120_ and BG505 Envs exhibited different conformations (Fig. 4c). Thus, after superimposing the gp120 residues in the BG24_LC-iGL_-GT1_N276gp120_ and BG24_mat_-BG505 structures, it was evident that the N276_gp120_ glycan in BG505 showed steric clashes with the CDRL1 iGL in BG24_LC-iGL_ (Fig. 4c). Flexibility of the N276_gp120_ glycan on BG505 may be more constrained than the counterpart glycan on GT1, as GT1 contains fewer N-glycans in the CD4bs, allowing for increased N276_gp120_ glycan flexibility. This assumption is consistent with the ELISA results showing that BG24_LC-iGL_ bound to GT1, but not to BG505 Env trimers with an N276_gp120_ glycan (Fig. 4a).

The only other known CD4bs-tageting bNAb with a helical CDRL1 is IOMA, another VH1-2*02 derived bNAb^27^. IOMA contains features that distinguish it from VRC01-class bNAbs, including a normal length (8 residue) CDRL3 and a 13-residue CDRL1, which adopts a short α-helix to accommodate the glycan at N276_gp120_. However, CLK31, an IOMA-like Ab isolated from naïve human B cells using a VRC01 germline-targeting immunogen, did not include a helical CDRL1^35^. Alignment of the LCs of BG24_LC-iGL_, IOMA, and CLK31 showed that each CDRL1 adopts a different configuration (Fig. 4d). These observations suggest CDRL1 helical conformations are diverse and have only been observed when bound to gp120s that contain the glycan at N276_gp120_.

### BG24_CDRL1-iGL_ accommodates the N276_gp120_ glycan in a non-engineered Env trimer

A longitudinal study that tracked the development of a VRC01-class lineage (PCIN63) found that bNAb development branched into two types of N276_gp120_ glycan engagement: one that interacted with and depended on the presence of the N276 _gp120_ glycan, and one in which CD4bs binding was diminished by the presence of the N276 _gp120_ glycan^32^. In the absence of longitudinal data for BG24_mat_ development, a BG24 intermediate, BG24_CDRL1-iGL_, was tested for neutralization against a 119-virus cross clade panel to better understand how the germline BG24 CDRL1 interacted with HIV-1 Envs bearing the N276 _gp120_ glycan. BG24_CDRL1-iGL_ exhibited neutralization activity against two viruses that contained PNGSs at N276_gp120_: clade D 6405.v4.c34 (6405) and clade CD 6952.v1.c20 (6952) (Fig. 5a). The 6405 Env was selected for further investigation by creating a soluble 6405 SOSIP.664 trimer. Sequence alignment of 6405 and BG505 gp120s showed the amino acid identity in the CD4bs and V4 loops differed by more than 50% between BG505 and 6405 Envs (Extended Data Fig. 4). The 6405 gp120 sequence included similar CD4bs PNGSs as BG505, except for the absence of a PNGS at position 363_gp120_ and an added PNGS at position 465_gp120_ (Extended Data Fig. 4). Binding of BG24 Fab constructs to 6405 was assessed by ELISA. Consistent with neutralization results (Fig. 5a), ELISAs showed that BG24_CDRL1-iGL_ and BG24_mat_ each bound the 6405 SOSIP, whereas BG24_LC-iGL_ bound 6405 to a lesser extent (Fig. 5b). BG24_iGL_s did not bind detectably to 6405 (Fig. 5b).

**Fig. 5.**
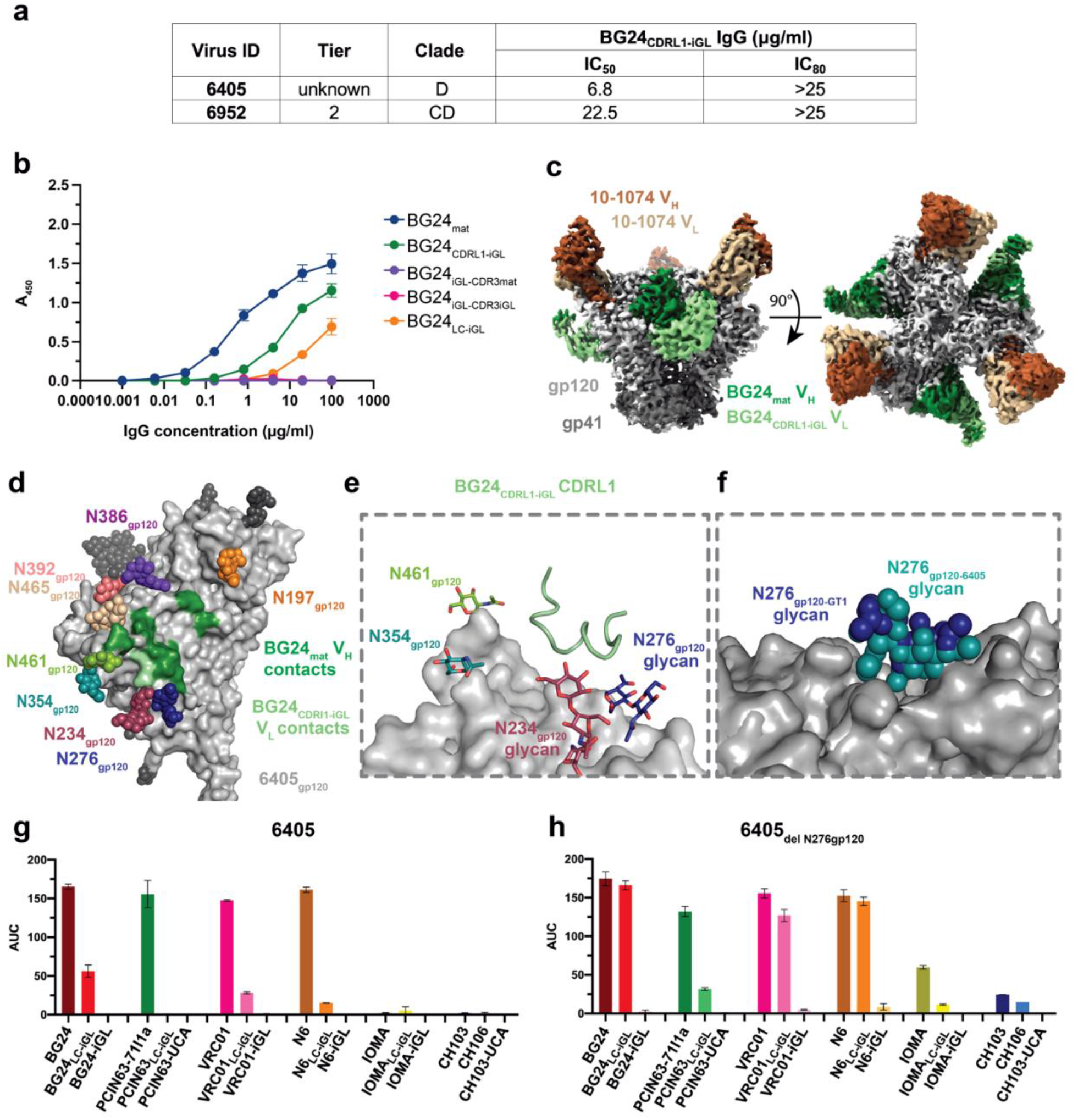
Non-engineered 6405 SOSIP recognizes BG24_CDRL1-iGL_. **a**, Summary of neutralization of 6405 and 6952 pseudoviruses by BG24_CDRL1-iGL_ IgGs. **b**, ELISA to access binding of BG24-derived Abs to 6405 SOSIP. Streptavidin plates were coated with randomly biotinylated SOSIPs and incubated with BG24-derived IgGs, at increasing concentrations. **c,** Side and top-down views of cryo-EM density of BG24_CDRL1-iGL_-6405-10-1074. **d**, Surface contacts made by BG24_CDRL1-iGL_ V_H_ (dark green) and V_L_ (light green) on 6405 gp120. **e**, Cartoon representation for the CDLR1 of BG24_CDRL1-iGL._ **f**, Alignment of GT1_N276gp120_ and 6405 gp120s in surface representation and N276 glycans in sphere representation. **g,h,** Summary for area under the curve (AUC) values derived from ELISAs that accessed binding of CD4bs IgGs to **g**, 6405 and **h,** 6405_delN276gp120_ SOSIPs. Streptavidin plates were coated with randomly biotinylated SOSIPs and incubated with CD4bs IgGs at increasing concentrations. Values are shown as mean of two individual replicates with associated error bars.

To understand how the germline CDRL1 of BG24 could be accommodated by a non-engineered Env trimer, we characterized interactions between BG24_CDRL1-iGL_ and 6405 SOSIP by solving a 3.4 Å cryo-EM structure of a BG24_CDRL1-iGL_-6405 complex (Fig. 5c, Extended Data Fig. 5a-d, Supplementary Table 1). As expected, BG24_CDRL1-iGL_ recognized the CD4bs of 6405, which contained N-glycans at positions N197_gp120_, N234_gp120_, N276_gp120_, N354_gp120_, N386_gp120_, N392_gp120_, N461_gp120_, and N465_gp120_ (Fig. 5d). Again, side chains were not modeled for CDRL1 residues (Extended Data Fig. 5e). As also observed in the BG24_LC-iGL_-GT1_N276_ structure, the CDRL1-iGL in the BG24_CDRL1-iGL_-6405 complex formed a helical conformation, although the CDRL1 confirmations in the two Fabs in these complexes were not identical (Extended Data Fig. 5f). By aligning gp120s from GT1_N276_ and 6405, we found that the well-ordered portions of the N276_gp120_ glycan occupied similar positions (Fig. 5f), suggesting that 6405 Env contains a native glycan landscape conducive to glycan flexibility at position N276_gp120_, similar to what was observed in this region of the GT1 Env structure.

We also evaluated binding of other CD4bs bNAbs to 6405 and a 6405_delN276gp120_ Env to determine if the CD4bs glycan landscape in 6405 was conducive to interactions with germline CDRL1s in other VRC01-class bNAbs (Fig. 5f-g). In this experiment, we included mature versions of the BG24, VRC01, N6 and IOMA bNAb Fabs, chimeric bNAb Fabs including an iGL LC, and complete iGL Fabs. PCIN63 and CH103 and intermediates were compared with their unmutated common ancestors (UCAs), as identified from longitudinal studies^32, 36^, instead of iGLs. The ELISA revealed that the 6405 Env interacted with VRC01_LC-iGL_ and N6_LC-iGL_ in addition to BG24_LC-iGL_. Binding for all species increased for 6405_del N276gp120_, indicating the N276_gp120_ glycan can sterically impede the CDRL1 iGL despite its flexibility. We conclude that the 6405 Env contains a CD4bs glycan landscape that tolerates the binding of germline VRC01-class CDRL1s.

## Discussion

VRC01-class bNAbs are promising targets for germline-targeting immunogen design as germline-encoded residues make signature contacts with gp120 that contribute to impressive breadth and potency^5, 18, 22, 23^. However, challenges in eliciting VRC01-class bNAbs through a germline-targeting vaccine regimen include explicitly selecting for the VH1-2*02 germline gene, overcoming CD4bs glycan barriers, and stimulating high levels of SHM^37, 38^. Overcoming these challenges will be facilitated by increased understanding of how germline VRC01-class Abs interact with germline-targeting immunogens.

Despite these challenges, progress has been made in developing a VRC01-class bNAb germline-targeting approach^21, 38^, which is initiated by engineering an immunogen that binds to the germline precursors of VRC01 bNAbs. Priming immunogens are engineered to interact with specific germline-encoded residues and lack CD4bs glycans that obstruct germline recognition^3, 29, 37, 39^. VRC01-class priming immunogens include monomeric gp120 cores^25, 37, 40^, SOSIP-based trimers^3^, and anti-idiotypic antibodies that recognize target BCRs with VH1-2*02 gene segments^41, 42^. Selecting a particular strain of Env and a gp120-versus trimeric Env-based platform to engineer priming characteristics is important. For example, a recent study comparing three different gp120-based immunogens found that the epitope and surrounding surface of the immunogen affects germline BCR selection in vivo^43^. Thus, identifying and developing the optimal priming immunogen for VRC01-like bNAb elicitation will require a robust understanding of the structural and biophysical nature of Env recognition by germline precursors.

In a sequential immunization approach, boosting immunogens are introduced to shape the development of a germline precursor into a bNAb by stimulating favorable SHMs^38^. Example boosting immunogens re-introduce native Env glycans and heterogenous Env strains to develop bNAbs capable of overcoming steric glycan barriers and have heterologous-neutralizing activity^38^. The N276_gp120_ glycan on HIV-1 Env provides a particularly difficult roadblock, as VRC01-like germline CDRL1s must become shorter or more flexible through SHM to avoid steric clashes_23,27,28,32,44_. Several iterations of this approach have been tested in animal models; however, the elicitation of heterologous neutralizing activity has not yet been accomplished^37, 45^.

BG24_mat_, represents a new VRC01-class bNAb that can be targeted for germline-targeting approaches^30^. BG24_mat_ has a fraction of the SHMs found in VRC01 and other VRC01-class bNAbs and maintains notable breadth and potency. Together with previous studies of the VRC01-class PCIN63 lineage and construction of a minimally mutated VRC01, our studies of BG24 suggest that high levels of SHM are not absolutely required for the development of VRC01-class Abs^32, 44^. Our cryo-EM structures of the iGL precursors of BG24 bound to the priming immunogen GT1 contribute to understanding how VRC01-class bNAb precursors interact with immunogens. We found that VH1-2*02 germline-encoded residues make the predicted signature contacts with gp120 and the long germline CDRL1 is accommodated in the absence of the N276_gp120_ glycan in GT1, rationalizing removal of this glycan in a priming immunogen since modeling suggested the germline CDRL1 conformation would clash with the N276_gp120_ glycan. These observations validate the design of priming immunogens that nurture interactions with germline residues and remove the N276_gp120_ glycan from the CD4bs epitope. We further investigated how the glycan landscape of an immunogen affects germline binding, finding that BG24_iGL-LC_ can evade clashes with the N276_gp120_ glycan when the BG24 HC includes bNAb features and the CD4bs epitope is only minimally glycosylated. Based on these observations, we propose that boosting immunogens might first aim to target mature HC features, and then introduce the N276_gp120_ glycan in a limited CD4bs glycan landscape before moving to fully glycosylated Env landscape.

We also characterized binding of BG24_CDRL1-iGL_ to the clade D 6405 Env, which suggested that some non-engineered HIV-1 Envs contain glycan landscapes that can accommodate germline VRC01-class CDRL1s. In the case of BG24_CDRL1-iGL_, accommodation of the N276_gp120_ glycan occurred through a helix-like conformation in the iGL CDRL1. Furthermore, ELISA data suggested that other VH1-2*02-derived bNAbs with iGL LCs can also bind to the 6405 Env.

Taken together, we believe the 6405 Env contains properties that make it a desirable candidate for a boosting immunogen to introduce the N276_gp120_ glycan after priming: in particular, unlike most CD4bs-targeting immunogens used as early boosting immunogens, 6405 is a non-engineered Env that can interact productively with germline LCs. Therefore, we propose further investigation of 6405 Env in boosting regimens to determine if these structural and biochemical observations translate in vivo.

## Methods

### BG24_iGL_ constructs design

Genes encoding the IGHV1-2*02 and IGLV2-11*02 germline sequences with mature CDR3 loops were used to generate the BG24_iGL-CDR3mat_ Fab construct. For the BG24_iGL-CDR3iGL_ construct, amino acids in D- and J-gene regions were reverted based on inferred sequences using IMGT.

### Protein expression and purification

Fabs and IgGs were expressed and purified as previously described^46^. Briefly, Fabs were expressed by transient transfection using the Expi293 expression system (ThermoFisher). Fab expression vectors contained genes of LC and the C-terminally 6x-His tagged HC. The Fab and IgG proteins were purified from cell supernatants by Ni^2+^-NTA (GE Healthcare) and protein A affinity chromatography (GE Healthcare), respectively, followed by size exclusion chromatography (SEC) using a Superdex 200 10/300 column (GE Healthcare).

SOSIP.664 Env constructs contained the disulfide mutations 501C and 650C (SOS), I55P (IP), and the furin cleavage site mutated to six arginine residues (6R)^33^. Genes encoding BG505 SOSIP.664v4.1-GT1 and 6405 SOSIPs were expressed using by transient transfection of Expi293 cells (ThermoFisher) and purified as described previously^47^. The 6405 SOSIP construct contained gp120 residues 46-477 from the 6405 sequence, with remaining gp120 residues derived from BG505 and the extracellular portion of the BG505 gp41^48^. Trimeric Env was separated from cell supernatants by PGT145 immunoaffinity chromatography, and SEC using a Superose 6 10/300 column (GE Healthcare) as described^49^.

### X-ray crystallography

Purified BG24_iGL-CDR3mat_ Fab was concentrated to 8-15 mg/mL. Matrix crystallization screens were performed at room temperature using the sitting drop vapor diffusion method by mixing equal volumes of protein sample and reservoir using a TTP LabTech Mosquito robot and commercially-available screens (Hampton Research and Qiagen). Initial hits were optimized and crystals were obtained in 20% PEG 3350 at 20 °C. Crystals were cryo-protected in glycerol stepwise until 20% before being cryopreserved in liquid nitrogen.

X-ray diffraction data were collected to 1.4 Å for BG24_iGL-CDR3mat_ Fab at the Stanford Synchroton Radiation Lightsource (SSRL) beamline 12-2 on a Pilatus 6M pixel detector (Dectris). Data from a single crystal were indexed and integrated in XDS^50^ and merged with AIMLESS in the CCP4 software suite^51^. Structures were determined by molecular replacement in PHASER^52^ using coordinates of the BG24_mat_ Fab (PDB 7UCE), after removal of CDR loops and independent searches of the V_H_V_L_ and C_H_C_L_ domains. Models were refined using rigid body and B-factor refinement in Phenix^53^, followed by several cycles of iterative manual building in Coot^54^ and real-space refinement with TLS groups in Phenix^53, 55^ (Supplementary Table 2).

### Assembly of protein complexes and cryo-EM sample preparation

Protein complexes for cryo-EM were generated by incubating a purified BG24_iGL_ Fab and the 10-1074 Fab with an Env trimer in a 3:1 Fab:trimer molar ratio and incubating at 4°C overnight. The complex was then SEC purified over a Superdex 200 1/150 column (GE Healthcare). The peak corresponding to complex was pooled and concentrated to 1.0 mg/ml. Quantifoil R2/2 400 mesh cryo-EM grids (Ted Pella) were prepared by glow-discharging for 1 min at 20 mA using a PELCO easiGLOW (Ted Pella). Fab-Env complexes (3 μL) were then applied to grids and blotted with Whatman No. 1 filter paper for 3-4 s at 100% humidity at room temperature. The grids were vitrified by and plunge-freezing in liquid ethane using a Mark IV Vitrobot (ThermoFisher).

### Cryo-EM data collection and processing

Data for single-particle cryo-EM were collected on either a Talos Artica (BG24_iGL-CDR3mat_-GT1-10-1074, BG24_iGL-CDR3iGL_ -GT1_N276gp120_ -10-1074, BG24_CDRL1-iGL_ -6405-10-1074) or a Titan Krios (BG24_iGL-CDR3mat_-GT1-10-1074) transmission electron microscope, operating at 200 kV and 300 kV, respectively. Movies were collected with beam-image shift over a single exposure per hole in a 3-by-3 pattern of 2 μm holes. For datasets collect on the Talos Artica, movies were recorded in super-resolution mode on a Falcon III camera (Thermo Fisher) at 1.436 Å•pixel^-1^ or a K3 camera (Gatan) at 0.4345 Å•pixel^-1^. Movies obtained from samples on the Titan Krios were collected in super-resolution mode on a K3 camera (Gatan) equipped with an BioQuantum energy filter (Gatan) with a 20 eV slit width at 0.4327 Å•pixel^-1^. The defocus range was set from 1.0-3.0 μm for each dataset.

The data processing workflow described below was preformed similarly for all datasets using RELION^56, 57^. Movies were motion-corrected using MotionCor2^58^ after binning. GCTF^59^ was used to estimate CTF, and micrograph power spectra that showed poor CTF fits or bad ice were removed. A subset of particles was manually picked and used for reference-free 2D classification. Classes representing the defined complex were used as references for RELION AutoPicking^56, 57^ to generate 2D classes. Subsequent 2D classes were inspected, and 2D classes representing a defined complex were selected for 3D classification. An ab initio model was generated using cryoSPARC^60^ using a subset of particles for each dataset and used as a reference in 3D classification which assumed C1 symmetry. 3D classes representing a defined complex were selected for 3D auto-refinement and post processing in Relion. Particles used in 3D refinement were then re-extracted and un-binned. Particles were then subjected to 3D classification with the map generated with un-binned particles used as a reference. Distinct classes representing a particular defined complex (C1 or C3 symmetric) were selected for 3D auto-refinement after masking out Fab C_H_C_L_ domains. Iterative rounds of particle CTF refinement, particle polishing, 3D auto-refinement, and post processing were used for each class to generate final maps. To improve resolution of Fab_LC_ CDRL1s, a soft mask surrounding the Fab VH-VL-gp120 interface was created in chimera and used for local refinements in cryoSPARC to improve density in this region and allow for CDRL1 fitting and refinement. Resolutions were calculated in RELION using the gold-standard FSC 0.143 criterion^61^. FSCs were generated by the 3DFSC program^62^.

### Cryo-EM model building and refinement

Model coordinates were generated by fitting reference gp120 (PDB 5T3Z), gp41(PDB 5T3Z), 10-1074 (PDB 5T3Z), and BG24-derivitive Fabs (this study) chains into cryo-EM density with UCSF Chimera^63^. Initial models were refined using the Phenix command *phenix.real_space_refine*^53, 55^. Sequence updates to the model and further manual refinement was conducted with Coot^54^. Iterative rounds of Phenix auto-refinement and manual refinements were done to generate the final models (Supplementary Table 1).

### Structural analyses

Structure figures were made using PyMol (Schrödinger LLC), UCSF Chimera^63^, and UCSF ChimeraX^64, 65^. PyMol was used to calculate r.m.s.d. values after pairwise alignment of Cα atoms. PDBePISA^66^ was used to calculate buried surface areas using a 1.4 Å probe. Calculations for gp120 BSA were for peptide components of gp120 and did not include glycan interactions. Defined interactions were assigned tentatively due to the low resolution of complexes using the following criteria: hydrogen bonds were assigned pairwise interactions that were less than 4.0 Å and with an A-D-H angle >90°, and van der Waals interactions were assigned as distances between atoms that were less than 4.0 Å.

### Enzyme-linked immunosorbent assay

SOSIP trimers were randomly biotinylated following manufacturer’s guidelines using the EZ-Link NHS-PEG4-Biotin kit (Thermo Fisher Scientific). The Pierce Biotin kit (Thermo Fisher Scientific) was used to quantify biotin molecules per SOSIP protomer: biotin estimations ranged from 1-10 biotin molecules per SOSIP protomers. Streptavidin-coated 96-well plates (Thermo Fisher Scientific) were coated with 5 µg/mL of randomly biotinylated SOSIPs diluted in 3% BSA in TBS-T (20mM Tris, 150 mM NaCl, 0.1% Tween20) and incubated at room temperature (RT) for 2 hours. Plates were washed to remove unbound SOSIPs. Serial dilutions of IgGs were made in 3% BSA in TBS-T and applied to the plates. After a 2-hour incubation at RT, plates were washed twice in TBS-T. Goat anti-human IgG Fc conjugated to horse-radish peroxidase (Southern BioTech,) was added at 1:8000 dilution for 30 minutes, followed by 3 washes with TBS-T. 1-Step™ Ultra TMB-ELISA Substrate Solution (ThermoFisher Scientific) was added for colorimetric detection and color development was quenched with 1N HCl. Absorbance was measured at 450nm. Two independent, biological replicates were performed.

## Acknowledgements

We thank J. Vielmetter, P. Hoffman, and the Protein Expression Center in the Beckman Institute at Caltech for expression assistance. Electron microscopy was performed in the Caltech Cryo-EM Center with assistance from S. Chen and A. Malyutin. We thank the Gordon and Betty Moore and Beckman Foundations for gifts to Caltech to support the Molecular Observatory (Dr. Jens Kaiser, Director) and the beamline staff at SSRL for data collection assistance. Use of the Stanford Synchrotron Radiation Lightsource, SLAC National Accelerator Laboratory, is supported by the U.S. Department of Energy, Office of Science, Office of Basic Energy Sciences under Contract No. DE-AC02-c76SF00515. The SSRL Structural Molecular Biology Program is supported by the DOE Office of Biological and Environmental Research, and by the National Institutes of Health, National Institute of General Medical Sciences (P41GM103393). The contents of this publication are solely the responsibility of the authors and do not necessarily represent the official views of NIGMS or NIH. C.O.B. was supported by a Burroughs Wellcome PDEP fellowship and HHMI Hanna Gray Fellowship. This work was supported by the National Institute of Allergy and Infectious Diseases (NIAID) Grant HIVRAD P01 AI100148 (to P.J.B. and M.C.N.), the Bill and Melinda Gates Foundation Collaboration for AIDS Vaccine Discovery (CAVD) grant INV-002143 (P.J.B., M.C.N.), and NIH P50 AI150464 (P.J.B.).

## Author contributions

K.A.D., C.O.B., H.B.G., T.S., M.C.N., and P.J.B. designed the research. K.A.D. performed experiments and K.A.D., C.O.B., and H.B.G. analyzed results. K.A.D. and P.J.B. wrote the manuscript with input from co-authors.

## Competing interests

The authors declare that there are no competing interests.

## Data availability

The atomic model generated for the X-ray crystallography structure of the BG24_iGL-CDR3mat_ Fab in this study has been deposited in the Protein Data Bank (PDB) under accession code 7UGM. The cryo-EM maps and atomic structures have been deposited in the PDB and/or Electron Microscopy Data Bank (EMDB) under accession codes 7UGN and EMD-26490 for BG24_iGL-CDR3iGL_-GT1-10-1074 Class 1, EMD-26491 for BG24_iGL-CDR3iGL_-GT1-10-1074 Class 2, 7UGO and EMD-26492 for BG24_iGL-CDR3mat_-GT1-10-1074, 7UGP and EMD-26493 forBG24_iGL-LC_-GT1_N276gp120_-10-1074 Class 1, EMD-26494 for BG24_iGL-LC_-GT1_N276gp120_-10-1074 Class 2, EMD-26495 for BG24_iGL-LC_-GT1_N276gp120_-10-1074 Class 3, and 7UGQ and EMD-26496 for BG24_CDRL-iGL_-6405-10-1074. Local refinement maps used to model CDRL1s of BG24-derivatives have been deposited with PDB and EMDB accession codes for each respective structure.

---

**Extended Data Fig. 1.**
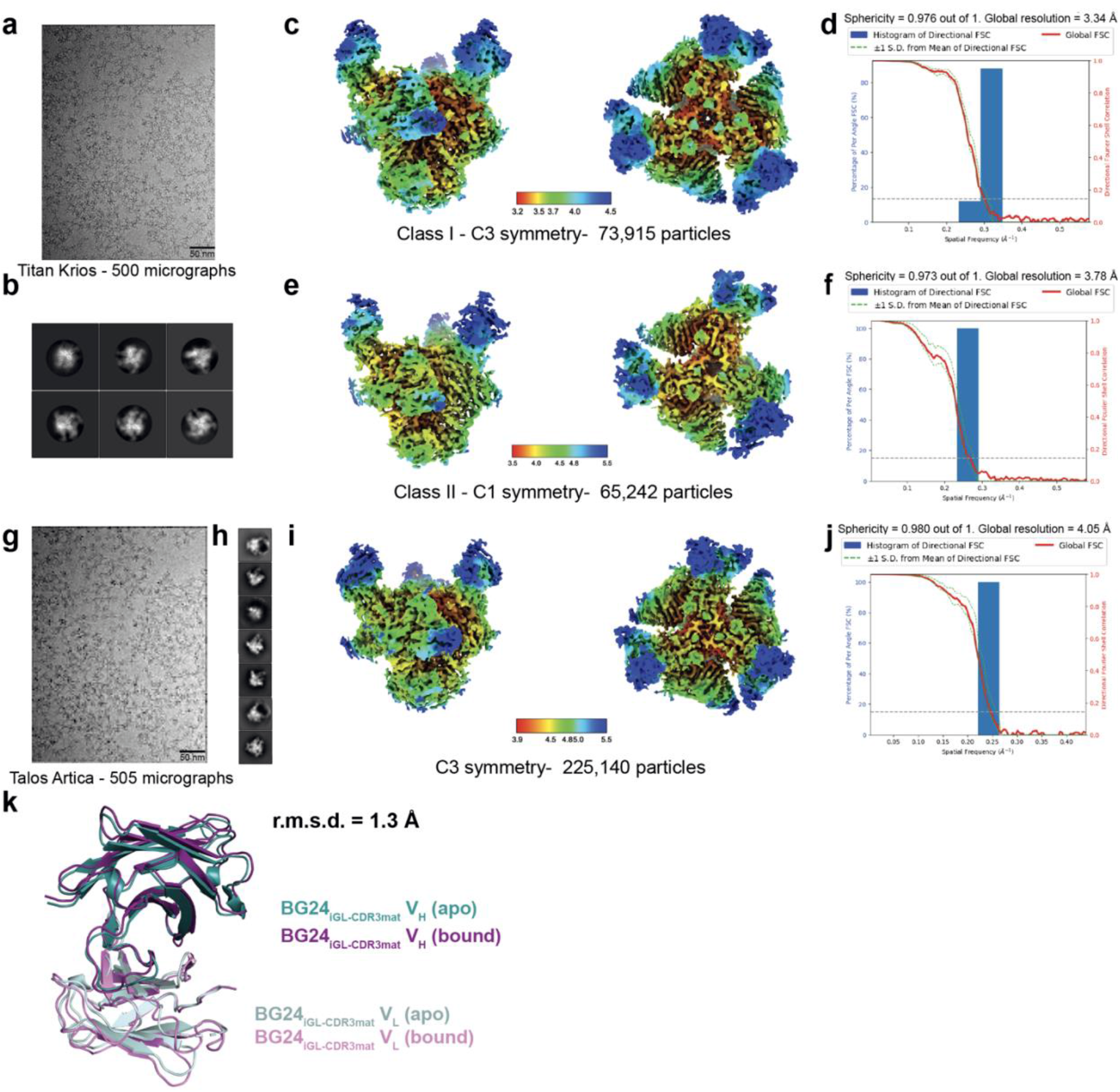
Cryo-EM data processing and validation for BG24_iGL_-GT1-10-1074 complexes and BG24_iGL-CDR3mat_ Fab alignment. **a-j**, Representative micrograph, cryo-EM 2D class averages, local resolution estimations, and gold-standard Fourier shell correlation (FSC) plots for **a-f**, BG24-iGL_CDR3iGL_-GT1-10-1074 and **g-j**, BG24_iGL-CDR3iGL_-GT1-10-1074. For the GT1-BG24_iGL-CDR3iGL_-10-1074 dataset, two classes were resolved: class I with three BG24_iGL-CDR3iGL_ Fabs bound to GT1, and class II with two BG24_iGL-CDR3mat_ Fabs bound to GT1. **k**, Alignment and r.m.s.d. (Å) of the crystal structure of apo BG24_iGL-CDR3mat_ Fab and the BG24_iGL-CDR3mat_ Fab bound to GT1 in the cryo-EM structure.

**Extended Data Fig. 2.**
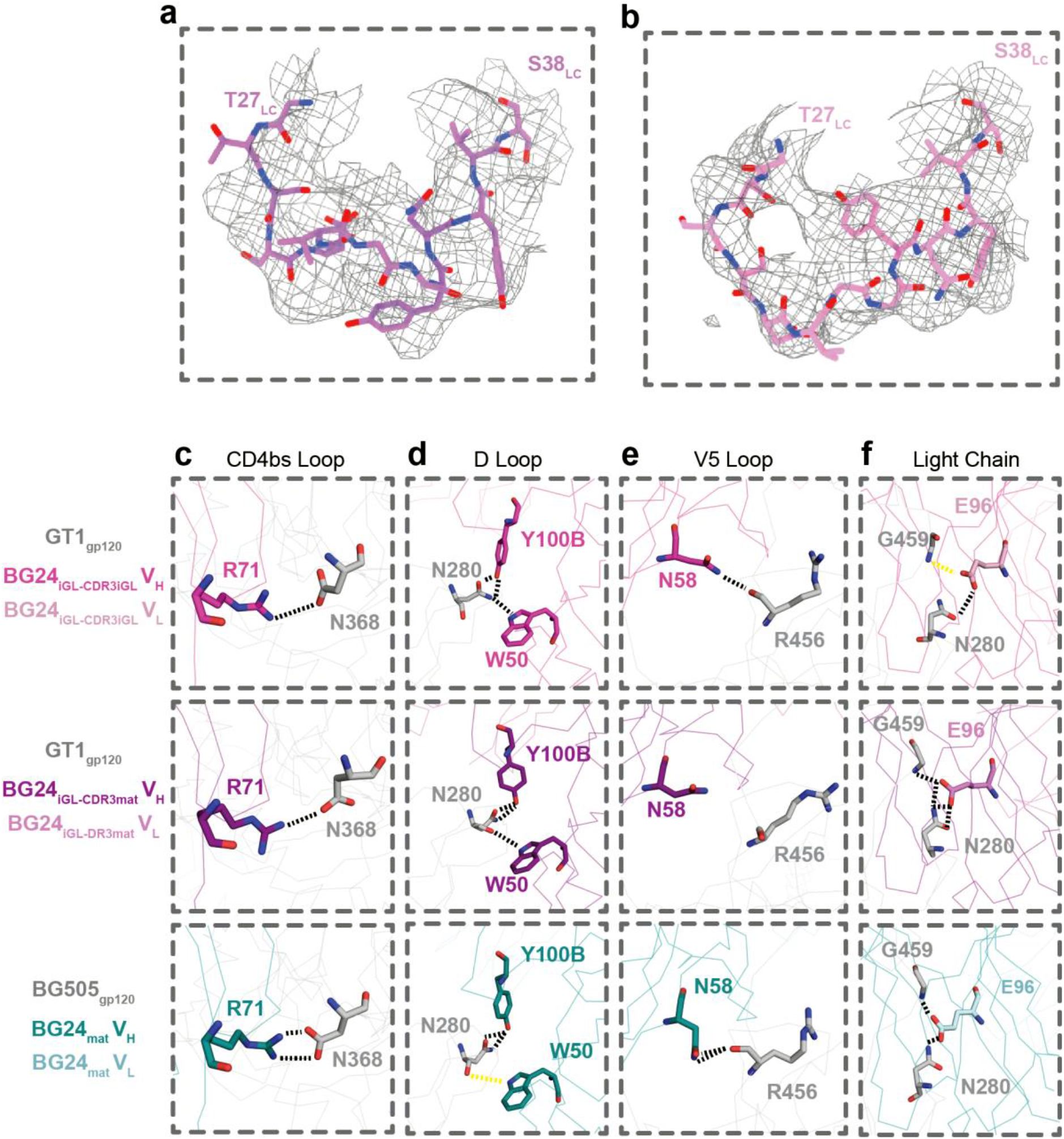
BG24_iGL_ CDRL1 local density and VH1-2*02 signature contacts comparison among BG24_iGL_ and BG24_mat_ structures. Local density for CDRL1 with modeled LC residues T27-S38_LC_ (stick representation) for **a**, BG24_iGL-CDR3mat_ and **b**, BG24_iGL-CDR3iGL_ contoured to 3.8 and 3.6 σ, respectively. Interactions in **c**, CD4bs, **d**, D Loop, **e**, V5 loop, and **f**, LC for BG24_iGL-CDR3iGL_-GT1_gp120,_ BG24_iGL-CDR3mat_-GT1_gp120_, and BG24_mat_-BG505_gp120_ structures. Contacts between atoms within 4 Å are represented by black dotted lines; contacts between 4-5 Å are represented by yellow dotted lines.

**Extended Data Fig. 3.**
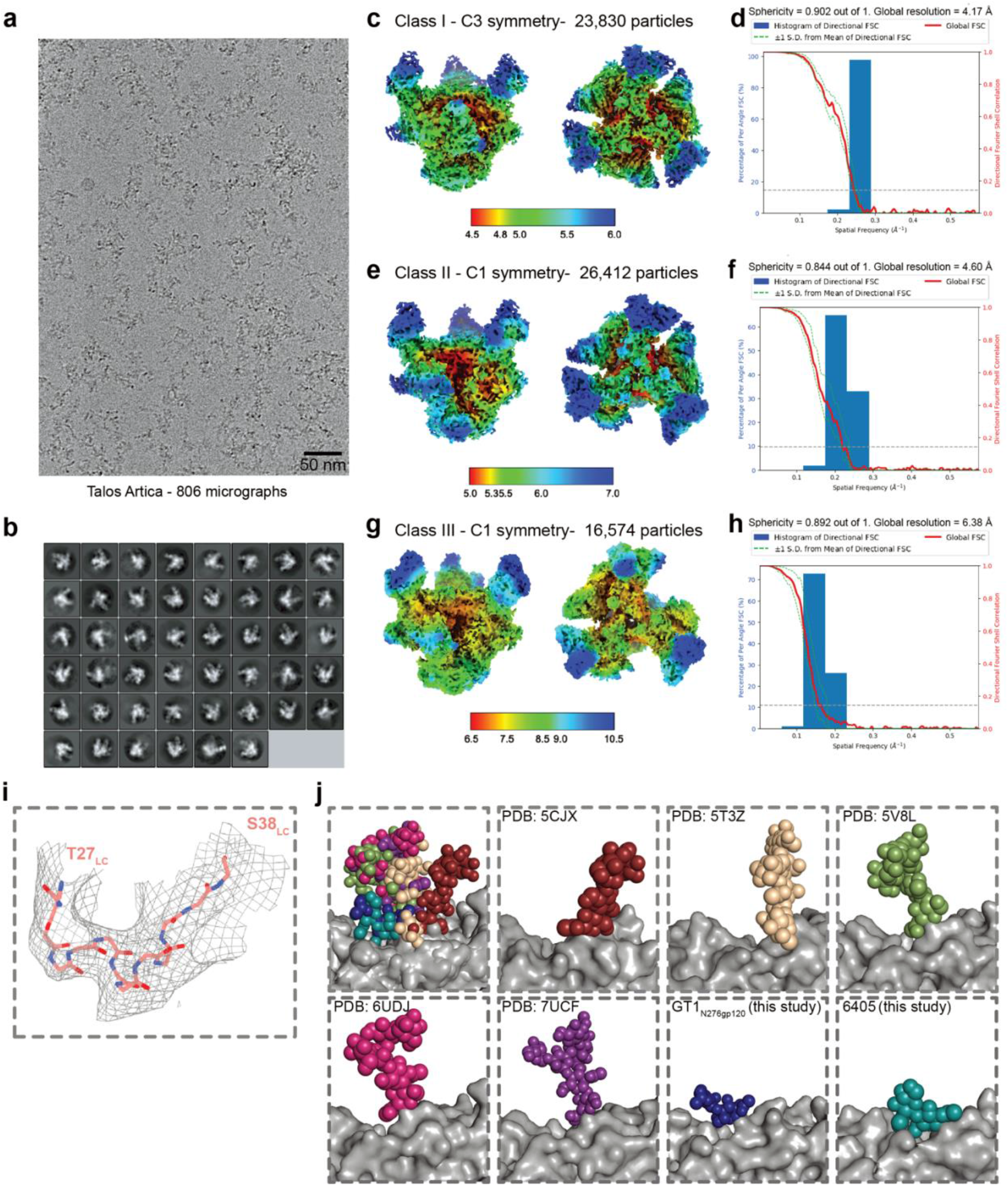
Cryo-EM data processing and validation for BG24_LC-iGL_-GT1_N276gp120_-10-1074 complex, BG24_LC-iGL_ CDRL1 local density, and analysis of N276_gp120_ glycan flexibility. **a-h**, Representative micrograph, cryo-EM 2D class averages, local resolution estimations, and gold-standard Fourier shell correlation (FSC) plots for BG24_LC-iGL_-GT1_N276gp120_-10-1074. For this dataset, three classes were resolved: class I with three BG24_LC-iGL_ Fabs bound to GT1_N276gp120_, class II with two BG24_LC-iGL_ Fabs bound to GT1_N276gp120_, and class III with one BG24_LC-iGL_ Fab bound to GT1_N276gp120_. **i**, Local density for CDRL1 with modeled backbone for LC residues T27-S38_LC_ (stick representation) for BG24_LC-iGL_ contoured to 5.4 σ. **j**, Comparison and overlay of the N276_gp120_ glycan from existing Env structures and GT1_N276gp120_ and 6405 from this study.

**Extended Data Fig. 4.**
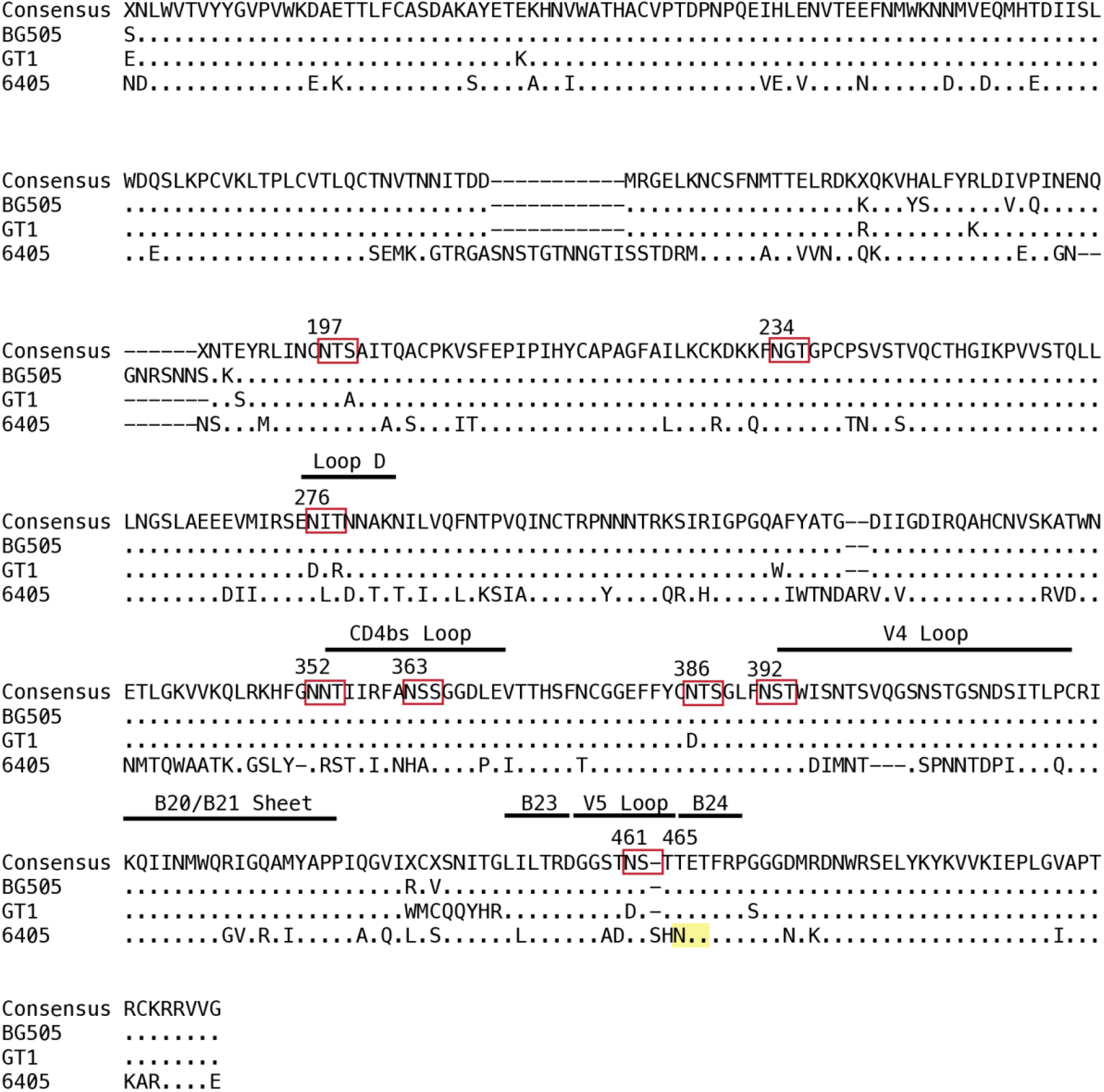
Sequence Alignment of BG505, GT1, and 6405 gp120. The sequence alignment for BG505, GT1, and 6405 gp120 residues and assigned consensus sequence. PNGS sequons in the consensus sequence are boxed in red, with residues numbers corresponding to Asn in the PNGS indicated above the sequence. PNGSs that are not conserved are highlighted in yellow, again with residues numbers corresponding to the Asn in the PNGS indicated above the sequence.

**Extended Data Fig. 5.**
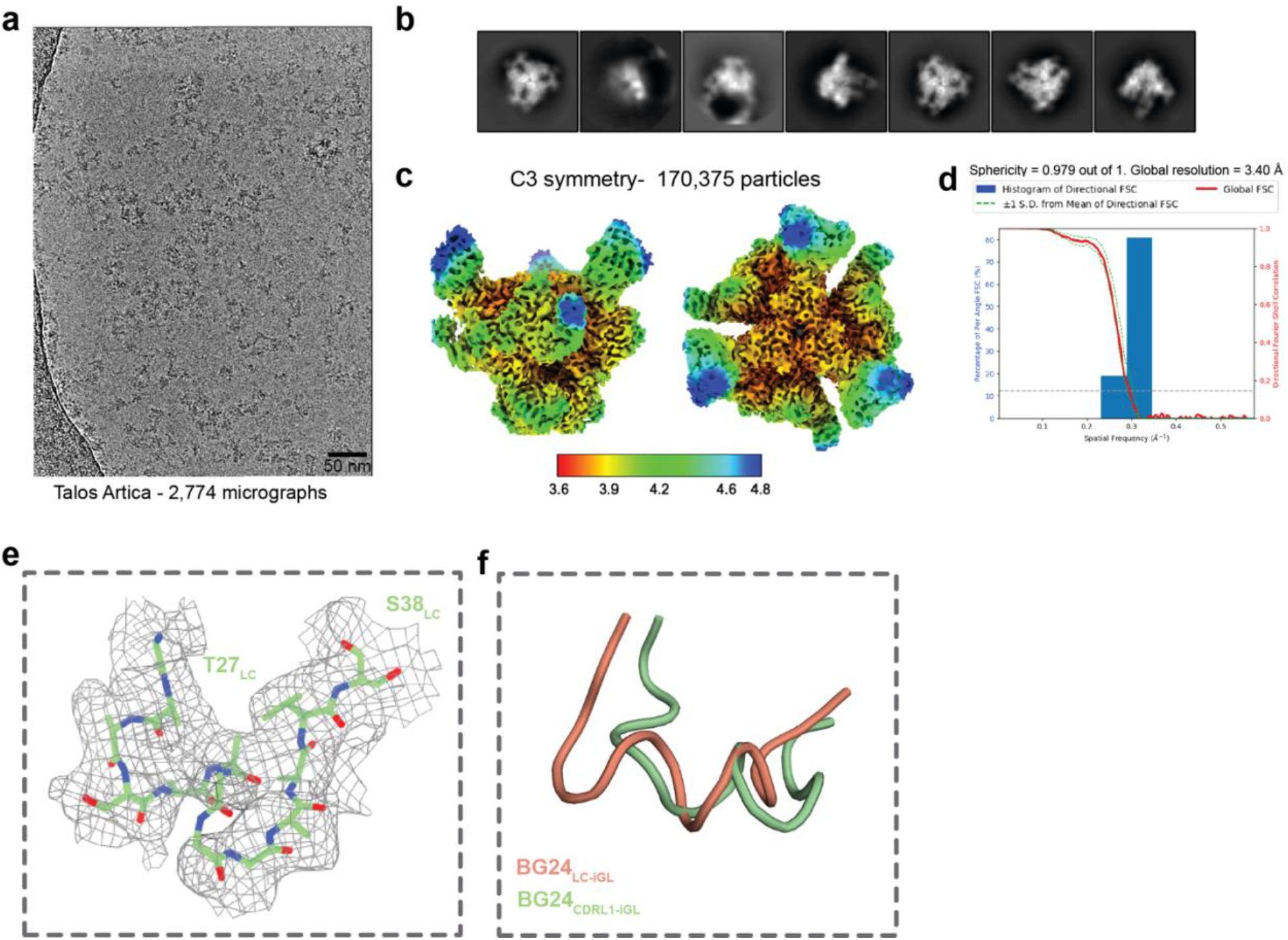
Cryo-EM data processing and validation for BG24_CDRL1-iGL_-6405-10-1074 complex and BG24_CDRL1-iGL_ CDRL1 local density. **a-d**, Representative micrograph, cryo-EM 2D class averages, local resolution estimations, and gold-standard Fourier shell correlation (FSC) plots for BG24_CDRL1-iGL_-6405-10-1074. **e**, Local density for CDRL1 with modeled backbone for residues T27-S38_LC_ (stick representation) for BG24_CDRL1-iGL_ contoured to 4.4 σ. **f**, Alignment of BG24_iGL_ CDLR1s from the BG24_LC-iGL_-GT1_N276gp120_-10-1074 structure and from the BG24_CDRL1-iGL_-6405-BG24_LC-iGL_-10-1074 structure. CDRL1s are represented in cartoon.

**Supplementary table 1.**
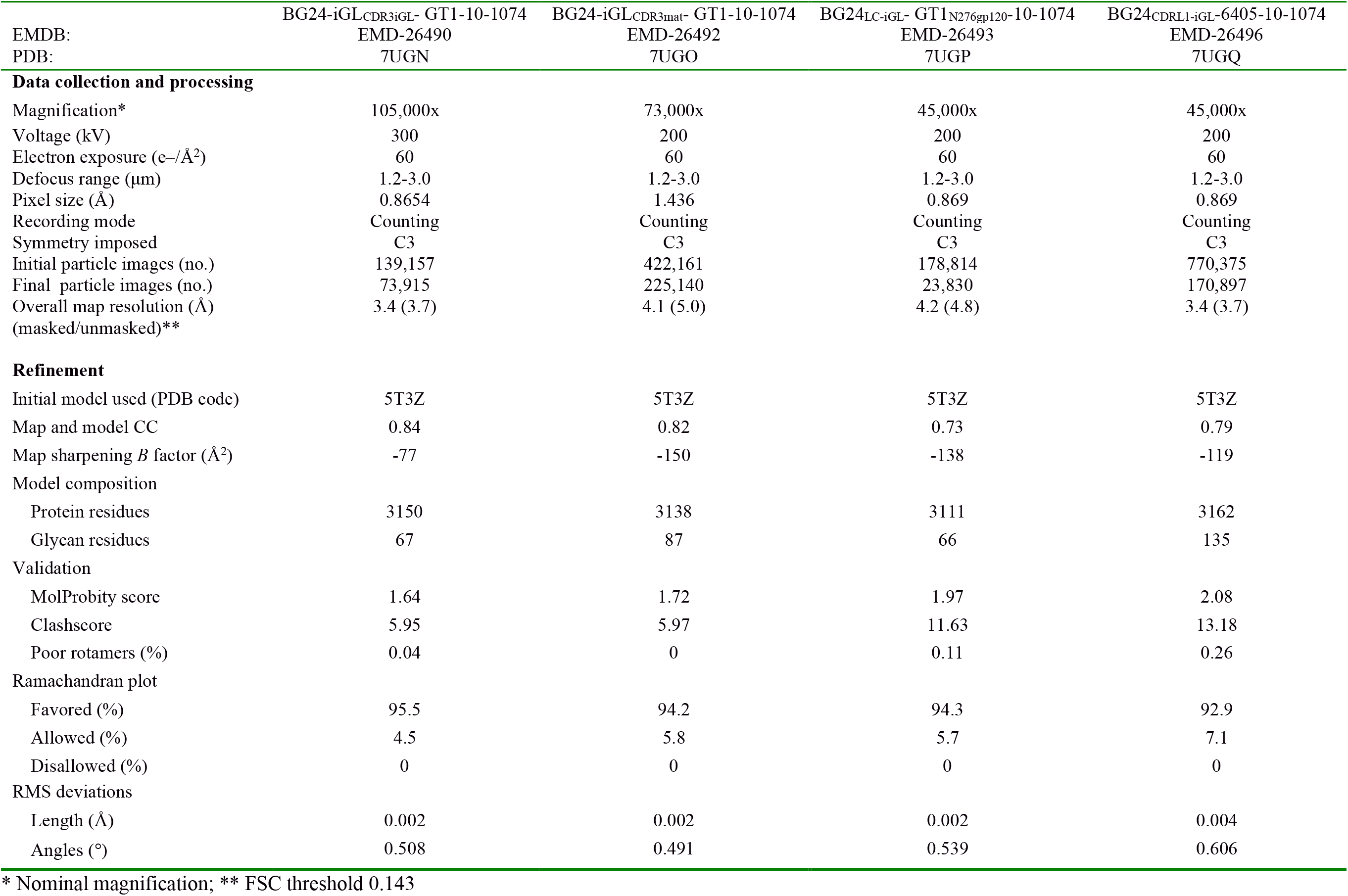
Cryo-EM data collection, refinement, and validation statistics.

**Supplementary table 2.** X-ray data collection and refinement statistics (molecular replacement)

| **PDB ID** | **iGL BG24 Fab**<br>**(12-2, SSRL)**<br>**7UGM** |
| --- | --- |
| **Data collection<sup>a</sup>** |  |
| Space group | P2<sub>1</sub>2<sub>1</sub>2<sub>1</sub> |
| Unit cell (Å) | 53.2, 70.8, 134.9 |
| $\alpha$, $\beta$, $\gamma$ (°) | 90, 90, 90 |
| Wavelength (Å) | 1.0 |
| Resolution (Å) | 38-1.4 (1.42-1.4) |
| Unique Reflections | 101,139 (49,320) |
| Completeness (%) | 100 (99.9) |
| Redundancy | 19.7 (19.2) |
| CC<sub>1/2</sub> (%) | 99.3 (67.2) |
| $\langle I/\sigma I \rangle$ | 11.0 (1.2) |
| Mosaicity (°) | 0.17 |
| R<sub>merge</sub> (%) | 16 (103) |
| R<sub>pim</sub> (%) | 3.7 (237) |
| Wilson *B*-factor | 14.2 |
| **Refinement and Validation** |  |
| Resolution (Å) | 38-1.4 |
| Number of atoms |  |
| Protein | 3,189 |
| Water | 447 |
| R<sub>work</sub>/R<sub>free</sub> (%) | 19.3/20.8 |
| R.m.s. deviations |  |
| Bond lengths (Å) | 0.006 |
| Bond angles (°) | 0.93 |
| MolProbity score | 1.42 |
| Clashscore (all atom) | 4.6 |
| Poor rotamers (%) | 0.3 |
| Ramachandran plot |  |
| Favored (%) | 96.9 |
| Allowed (%) | 3.1 |
| Disallowed (%) | 0.2 |
| Average *B*-factor (Å) |  |
| Protein | 24.3 |
| Water | 32.7 |

## Notes

### Competing Interest Statement

The authors have declared no competing interest.

## References

1. Escolano, A. et al. Immunization expands B cells specific to HIV-1 V3 glycan in mice and macaques. Nature 570, 468–473 (2019).

2. Lee, J. H. & Crotty, S. HIV vaccinology: 2021 update. Semin. Immunol. 51, 101470 (2021).

3. Medina-Ramírez, M. et al. Design and crystal structure of a native-like HIV-1 envelope trimer that engages multiple broadly neutralizing antibody precursors in vivo. J. Exp. Med. 214, 2573–2590 (2017).

4. Sanders, R. W. & Moore, J. P. Native-like Env trimers as a platform for HIV-1 vaccine design. Immunol. Rev. 275, 161–182 (2017).

5. Zhou, T. Structural Basis for Broad and Potent Neutralization of HIV-1 by Antibody VRC01. 1–8 (2010).

6. Schommers, P. et al. Restriction of HIV-1 Escape by a Highly Broad and Potent Neutralizing Antibody. Cell 180, 471–489.e22 (2020).

7. Huang, J. et al. Identification of a CD4-Binding-Site Antibody to HIV that Evolved Near-Pan Neutralization Breadth. Immunity 45, 1108–1121 (2016).

8. Scheid, J. F. et al. Sequence and Structural Convergence of Broad and Potent HIV Antibodies That Mimic CD4 Binding. Science 333, 1633–1637 (2011).

9. Wu, X. et al. Rational Design of Envelope Identifies Broadly Neutralizing Human Monoclonal Antibodies to HIV-1. Science 329, 856–861 (2010).

10. Bar, K. J. et al. Effect of HIV Antibody VRC01 on Viral Rebound after Treatment Interruption. N. Engl. J. Med. 375, 2037–2050 (2016).

11. Bar-On, Y. et al. Safety and antiviral activity of combination HIV-1 broadly neutralizing antibodies in viremic individuals. Nat. Med. 24, 1701–1707 (2018).

12. Miner, M. D., Corey, L. & Montefiori, D. Broadly neutralizing monoclonal antibodies for HIV prevention. J. Int. AIDS Soc. 24, (2021).

13. Mendoza, P. et al. Combination therapy with anti-HIV-1 antibodies maintains viral suppression. Nature 1–21 (2018) doi:10.1038/s41586-018-0531-2.

14. VRC 602 Study Team et al. Safety, pharmacokinetics and neutralization of the broadly neutralizing HIV-1 human monoclonal antibody VRC01 in healthy adults. Clin. Exp. Immunol. 182, 289–301 (2015).

15. Lynch, R. M. et al. Virologic effects of broadly neutralizing antibody VRC01 administration during chronic HIV-1 infection. Sci. Transl. Med. 7, (2015).

16. Mayer, K. H. et al. Safety, pharmacokinetics, and immunological activities of multiple intravenous or subcutaneous doses of an anti-HIV monoclonal antibody, VRC01, administered to HIV-uninfected adults: Results of a phase 1 randomized trial. PLOS Med. 14, e1002435 (2017).

17. Escolano, A., Dosenovic, P. & Nussenzweig, M. C. Progress toward active or passive HIV-1 vaccination. J. Exp. Med. 214, 3–16 (2017).

18. West Jr, A. P. Structural basis for germ-line gene usage of a potent class of antibodies targeting the CD4-binding site of HIV-1 gp120. 1–8 (2012) doi:10.1073/pnas.1208984109/-/DCSupplemental.

19. Xiao, X. et al. Germline-like predecessors of broadly neutralizing antibodies lack measurable binding to HIV-1 envelope glycoproteins: Implications for evasion of immune responses and design of vaccine immunogens. Biochem. Biophys. Res. Commun. 390, 404–409 (2009).

20. Bonsignori, M. et al. Analysis of a Clonal Lineage of HIV-1 Envelope V2/V3 Conformational Epitope-Specific Broadly Neutralizing Antibodies and Their Inferred Unmutated Common Ancestors. J. Virol. 85, 9998–10009 (2011).

21. Burton, D. R. Advancing an HIV vaccine; advancing vaccinology. Nat. Rev. Immunol. 19, 77–78 (2019).

22. Borst, A. Germline VRC01 antibody recognition of a modified clade C HIV-1 envelope trimer and a glycosylated HIV-1 gp120 core. 1–32 (2018).

23. Scharf, L. et al. Structural basis for HIV-1 gp120 recognition by a germ-line version of a broadly neutralizing antibody. Proc. Natl. Acad. Sci. 110, 6049–6054 (2013).

24. Zhou, T. et al. Structural Repertoire of HIV-1-Neutralizing Antibodies Targeting the CD4 Supersite in 14 Donors. Cell 161, 1280–1292 (2015).

25. Jardine, J. G. et al. HIV-1 broadly neutralizing antibody precursor B cells revealed by germline-targeting immunogen. science 351, 1458–1463 (2016).

26. Zhou, T. et al. Quantification of the Impact of the HIV-1-Glycan Shield on Antibody Elicitation. Cell Rep. 19, 719–732 (2017).

27. Gristick, H. B. et al. Natively glycosylated HIV-1 Env structure reveals new mode for antibody recognition of the CD4-binding site. Nat. Publ. Group 23, 906–915 (2016).

28. Zhou, T. et al. Multidonor Analysis Reveals Structural Elements, Genetic Determinants, and Maturation Pathway for HIV-1 Neutralization by VRC01-Class Antibodies. Immunity 39, 245–258 (2013).

29. Jardine, J. G. et al. Priming a broadly neutralizing antibody response to HIV-1 using a germline-targeting immunogen. Science 349, 156–161 (2015).

30. Barnes, C. O. et al. A naturally arising broad and potent CD4-binding site antibody with low somatic mutation. BioRxiv 51 (2022) doi:https://www.biorxiv.org/content/10.1101/2022.03.16.484662v1.

31. Huang, D. et al. B cells expressing authentic naive human VRC01-class BCRs can be recruited to germinal centers and affinity mature in multiple independent mouse models. Proc. Natl. Acad. Sci. 117, 22920–22931 (2020).

32. Umotoy, J. et al. Rapid and Focused Maturation of a VRC01-Class HIV Broadly Neutralizing Antibody Lineage Involves Both Binding and Accommodation of the N276-Glycan. Immunity 51, 141–154.e6 (2019).

33. Sanders, R. W. et al. A Next-Generation Cleaved, Soluble HIV-1 Env Trimer, BG505 SOSIP.664 gp140, Expresses Multiple Epitopes for Broadly Neutralizing but Not Non-Neutralizing Antibodies. PLoS Pathog. 9, e1003618-20 (2013).

34. Mouquet, H. et al. Complex-type N-glycan recognition by potent broadly neutralizing HIV antibodies. Proc. Natl. Acad. Sci. 109, E3268–E3277 (2012).

35. Havenar-Daughton, C. The human naive B cell repertoire contains distinct subclasses for a germline-targeting HIV-1 vaccine immunogen. 1–15 (2018).

36. Liao, H.-X. et al. Co-evolution of a broadly neutralizing HIV-1 antibody and founder virus. Nature 496, 469–476 (2013).

37. Parks, K. R. et al. Overcoming Steric Restrictions of VRC01 HIV-1 Neutralizing Antibodies through Immunization. CellReports 29, 3060–3072.e7 (2019).

38. Stamatatos, L., Pancera, M. & McGuire, A. T. Germline-targeting immunogens. *Immunol. Rev.* **275**, 203–216 (2017).

39. McGuire, A. T. et al. Specifically modified Env immunogens activate B-cell precursors of broadly neutralizing HIV-1 antibodies in transgenic mice. Nat. Commun. 7, 10618 (2016).

40. Jardine, J. et al. Rational HIV Immunogen Design to Target Specific Germline B Cell Receptors. Science 340, 711–716 (2013).

41. Seydoux, E. et al. Development of a VRC01-class germline targeting immunogen derived from anti-idiotypic antibodies. Cell Rep. 35, 109084 (2021).

42. Dosenovic, P. et al. Anti-idiotypic antibodies elicit anti-HIV-1–specific B cell responses. J. Exp. Med. 216, 2316–2330 (2019).

43. Lin, Y.-R. et al. HIV-1 VRC01 Germline-Targeting Immunogens Select Distinct Epitope-Specific B Cell Receptors. Immunity 53, 840–851.e6 (2020).

44. Jardine, J. G. et al. Minimally Mutated HIV-1 Broadly Neutralizing Antibodies to Guide Reductionist Vaccine Design. PLoS Pathog. 12, e1005815–33 (2016).

45. Briney, B. et al. Tailored Immunogens Direct Affinity Maturation toward HIV Neutralizing Antibodies. Cell 166, 1459–1470.e11 (2016).

46. Yang, Z., Wang, H., Liu, A. Z., Gristick, H. B. & Bjorkman, P. J. Asymmetric opening of HIV-1 Env bound to CD4 and a coreceptor-mimicking antibody. Nat. Struct. Mol. Biol. 26, 1167–1175 (2019).

47. Escolano, A. et al. Sequential immunization of macaques elicits heterologous neutralizing antibodies targeting the V3-glycan patch of HIV-1 Env. Sci. Transl. Med. 13, eabk1533 (2021).

48. Joyce, M. G. et al. Soluble Prefusion Closed DS-SOSIP.664-Env Trimers of Diverse HIV-1 Strains. CellReports 21, 2992–3002 (2017).

49. Cupo, A. et al. Optimizing the production and affinity purification of HIV-1 envelope glycoprotein SOSIP trimers from transiently transfected CHO cells. PLOS ONE 14, e0215106 (2019).

50. Kabsch, W. Integration, scaling, space-group assignment and post-refinement. Acta Crystallogr. D Biol. Crystallogr. 66, 133–144 (2010).

51. Winn, M. D. et al. Overview of the *CCP* 4 suite and current developments. Acta Crystallogr. D Biol. Crystallogr. 67, 235–242 (2011).

52. McCoy, A. J. et al. Phaser crystallographic software. J. Appl. Crystallogr. **40**, 658– 674 (2007).

53. Adams, P. D. et al. *PHENIX* : a comprehensive Python-based system for macromolecular structure solution. Acta Crystallogr. D Biol. Crystallogr. 66, 213–221 (2010).

54. Emsley, P., Lohkamp, B., Scott, W. G. & Cowtan, K. Features and development of *Coot*. Acta Crystallogr. D Biol. Crystallogr. **66**, 486–501 (2010).

55. Afonine, P. V. et al. Real-space refinement in *PHENIX* for cryo-EM and crystallography. Acta Crystallogr. Sect. Struct. Biol. 74, 531–544 (2018).

56. Zivanov, J. et al. New tools for automated high-resolution cryo-EM structure determination in RELION-3. eLife 7, e42166 (2018).

57. Scheres, S. H. W. RELION: Implementation of a Bayesian approach to cryo-EM structure determination. J. Struct. Biol. 180, 519–530 (2012).

58. Zheng, S. Q. et al. MotionCor2: anisotropic correction of beam-induced motion for improved cryo-electron microscopy. Nat. Methods 14, 331–332 (2017).

59. Zhang, K. Gctf: Real-time CTF determination and correction. J. Struct. Biol. 193, 1– 12 (2016).

60. Punjani, A., Rubinstein, J. L., Fleet, D. J. & Brubaker, M. A. cryoSPARC: algorithms for rapid unsupervised cryo-EM structure determination. Nat. Methods 14, 290–296 (2017).

61. Scheres, S. H. W. & Chen, S. Prevention of overfitting in cryo-EM structure determination. Nat. Methods 9, 853–854 (2012).

62. Tan, Y. Z. et al. Addressing preferred specimen orientation in single-particle cryo-EM through tilting. Nat. Methods 14, 793–796 (2017).

63. Pettersen, E. F. et al. UCSF Chimera?A visualization system for exploratory research and analysis. J. Comput. Chem. 25, 1605–1612 (2004).

64. Goddard, T. D. et al. UCSF ChimeraX: Meeting modern challenges in visualization and analysis: UCSF ChimeraX Visualization System. Protein Sci. 27, 14–25 (2018).

65. Pettersen, E. F. et al. UCSF CHIMERAX : Structure visualization for researchers, educators, and developers. Protein Sci. 30, 70–82 (2021).

66. Krissinel, E. & Henrick, K. Inference of Macromolecular Assemblies from Crystalline State. J. Mol. Biol. 372, 774–797 (2007).

